# Single-Cell Raman Profiling Enables Rapid Precision Phage Therapy Against Multidrug-Resistant Hypervirulent *Klebsiella pneumoniae*

**DOI:** 10.64898/2026.01.25.701629

**Authors:** Beimin Liu, Chao Wang, Qingyang Zeng, Min Wei, Weilai Lu, Xueyan Gao, Jing Wan, Jie Feng, Yu Vincent Fu

## Abstract

Multidrug-resistant hypervirulent *Klebsiella pneumoniae* (MDR-hvKP) poses a severe global health threat. Phage therapy is a promising alternative, but requires precise matching of phage to the bacterial strain. Here, we present a proof-of-concept method that integrates single-cell Raman spectroscopy with deep learning to enable rapid and precise selection of lytic phages against MDR-hvKP. By profiling Raman signatures of strains across multiple KL-types (capsule locus types), we trained three deep learning architectures for phage-host matching. Among them, the CNN_MLP-Transformer achieved the best performance (99.7%), slightly outperforming CNN_MLP (99.2%) and CNN_MLP-Attention (99.5%). Validation using 10 hvKP clinical isolates yielded an average phage selection accuracy of 78.3%. These findings demonstrate the feasibility and clinical potential of AI- augmented Raman spectroscopy as a rapid, label-free, precise strategy for guiding phage therapy against MDR-hvKP infections.

**Teaser:** AI-guided single-cell Raman profiling enables rapid precision phage selection against multidrug-resistant *K. pneumoniae*.

## INTRODUCTION

*Klebsiella pneumoniae* is a gram-negative, encapsulated, facultative anaerobic bacterium, which is commonly found in the normal flora of human mouth, skin, and intestines (*1*). Despite its status as a commensal organism, *K. pneumoniae* is an opportunistic pathogen capable of causing a range of severe infections, including pneumonia, recurrent urinary tract infections, and pyogenic liver abscesses (*2*). Alarmingly, it now accounts for nearly one-third of all Gram-negative bacterial infections worldwide. Moreover, *K. pneumoniae* is one of the ESKAPE pathogens, notorious for its ability to rapidly acquire multidrug resistance in clinical settings, making it a critical public health concern (*3*).

The emergence of hypervirulent *K. pneumoniae* (hvKP) in recent decades has further complicated public health threat. Unlike classical *K. pneumoniae* (cKP), which predominantly affects immunocompromised hosts, hvKP can cause severe invasive infections in healthy individuals that develop quickly and spread to various body sites with high morbidity and mortality. The convergence of hypervirulence and multidrug resistance has led to the rise of multidrug-resistant hypervirulent strains (MDR-hvKP), which pose an intractable clinical challenge (*4, 5*). In 2024, the World Health Organization warned countries about increasing reports of MDR-hvKP. This issue is further exacerbated by the stagnation of antibiotic development since the 1980s, underscoring the urgent need for alternative antimicrobial strategies (*6*).

Bacteriophage therapy offers a promising solution. Bacteriophages are viruses that specifically infect bacteria. Lytic phages, in particular, can rapidly kill target bacteria, offering a potent therapeutic means against antibiotic-resistant infections (*7*). Recent studies have demonstrated the efficacy of phage therapy against drug-resistant pathogens such as *Pseudomonas aeruginosa*, *Staphylococcus aureus*, and *Acinetobacter baumannii (8-12)*. In the case of *K. pneumoniae*, encouraging results have been reported in murine infection models and in clinical settings, including treatment of recurrent urinary tract infections and the clearance of *K. pneumoniae* biofilms in prosthetic joint infections (*13-15*).

However, a major limitation of phage therapy lies in the narrow host range of most phages. In many cases, a phage can infect only a single *K. pneumoniae* strain. Effective treatment therefore requires accurate and rapid matching between phages and the infecting strain (*16*). Two primary strategies have been proposed to address this challenge. The first involves the use of broad-spectrum phage cocktails designed to lyse a wide array of bacterial strains. However, the diversity of bacterial strains makes comprehensive cocktail formulation extremely difficult to tailor. The second approach relies on targeted phage selection guided by bacterial typing. This targeted strategy, while more precise, depends on the availability of rapid, reliable, and cost-effective diagnostic methods for bacterial typing, as well as a comprehensive phage library.

Phage-bacterium interactions are typically initiated through recognition and binding to specific surface structures such as porins, pili, lipopolysaccharides (O-antigen), and capsular polysaccharides (K-antigen) (*17, 18*). The capsule, which constitutes the outermost layer of *K. pneumoniae*, often hinders phage binding unless the phage encodes depolymerases to degrade the polysaccharide capsule (*19-21*). These depolymerases exhibit exceptional specificity, and usually target only a single capsule type of *K. pneumoniae*. Consequently, phage infectivity is largely dependent on capsule type specificity of *K. pneumoniae* strains (*22, 23*).

More than 79 distinct capsule serotypes (K-type) have been described in *K. pneumoniae*, each with unique capsular polysaccharide features that influence phage susceptibility (*24-26*). However, capsular serotyping of *K. pneumoniae* is seldom performed in clinical laboratories due to methodological complexity and cost constraints. The classical Quellung reaction (or Neufeld test), which relies on K-type specific antisera to induce capsular swelling, are labor-intensive, subjective, and costly, and their utility is limited by antisera availability. Alternative immunochromatographic strip assay offers faster workflows but suffers from restricted serotype coverage and substantial false-negative rates (*27, 28*).

With the advent of next-generation DNA sequencing, capsule classification of *K. pneumoniae* has shifted from phenotypic K-serotyping to genotypic characterization of the capsular polysaccharide synthesis (CPS) locus (*29*). Capsule types were further designated as KL-types (capsule locus types), each representing a distinct genetic architecture within the *cps* cluster such as variations in *wzx*, *wzy*, glycosyltransferases, sugar-synthesis, and modification enzymes genes (*30, 31*). Subtle changes in gene clusters lead to significant variations in capsular composition and phage sensitivity consequently. Accordingly, our therapeutic phage library is organized according to KL-type specific infectivity profiles of phages.

To date, more than 158 KL-types have been described across *K. pneumoniae* populations (*32*). However, the routine determination of KL-type in clinical settings remains challenging. Accurate identification of KL-type requires whole-genome sequencing with high-quality *cps*-locus assemblies. The *cps* locus is highly variable in size, composition, and repetitive elements, making it difficult to assemble from short-read sequencing data, and incomplete assemblies often lead to false KL-type calls (*29*). Furthermore, DNA sequencing is costly, time-consuming, and cannot reliably resolve KL-type from small bacterial loads, which is an inherent limitation when analyzing early-stage or low-abundance clinical samples.

Raman spectroscopy has recently emerged as a promising tool for *K. pneumoniae* capsular serotyping (*33*). By measuring the inelastic photon scattering associated with molecular vibrations of chemical bonds, Raman spectroscopy provides rich biochemical fingerprints of individual cells. Subtle differences in capsular polysaccharide composition among various serotypes may yield distinct spectral patterns. Meanwhile, Raman spectroscopy is rapid, label-free, highly specific, non-destructive, and minimally affected by water, enabling direct analysis of live microbes in complex clinical samples such as blood, urine, and pus, with minimal preparation (*34-36*). While the complexity of Raman spectra has historically limited their interpretability, advances in artificial intelligence (AI), particularly deep learning, now enable robust extraction of biologically meaningful features from high-dimensional spectral data (*37-39*). The integration of Raman spectroscopy with deep learning has already shown promise in microbial species identification, antibiotic susceptibility testing, and metabolic phenotyping (*36, 40-43*).

Here, we present a proof-of-concept method that combines single-cell Raman spectroscopy with deep learning for rapid phage selection against MDR-hvKP. We acquired Raman spectra from 42 MDR-hvKP strains encompassing 17 KL-types and trained three deep learning models: CNN_MLP (a convolutional neural network combined with a multilayer perceptron), CNN_MLP-Attention, and CNN_MLP-Transformer. The CNN_MLP-Transformer model outperformed the others, achieving a phage match accuracy of 99.7%, compared to both CNN_MLP and CNN_MLP-Attention. Validation of 10 independent clinical isolates confirmed the feasibility of this method, supporting its potential utility for guiding precise phage therapy against MDR-hvKP infections.

## RESULT

### Construction of the MDR-hvKP Raman spectral library

Given the pivotal role of capsular polysaccharide in phage recognition, our therapeutic phage library was systemically organized based on the KL-type specific lytic activity of each phage. Once the KL-type of a clinical isolate is determined, the corresponding lytic phage can be rapidly and rationally selected for treatment. As a proof of concept for establishing a clinically practical phage-selection framework, we prioritized 17 representative KL-types of *K. pneumoniae*, which account for the vast majority of multidrug-resistant and hypervirulent *K. pneumoniae* infections worldwide (*26, 44*).

To construct a training spectral dataset for the deep learning, we collected 42 MDR-hvKP strains representing these 17 KL-types based on whole genome sequencing. In total, 6,254 single-cell Raman spectra were acquired (table S1). For less prevalent KL-types, such as KL20, KL51 and KL62, we acquired additional spectra per strain to ensure a balanced dataset. Fig. 1A presents the averaged Raman spectra for the 17 KL-types, and the average Raman spectra of the 42 strains used for model training are shown in Fig. 1B.

**Fig. 1.**
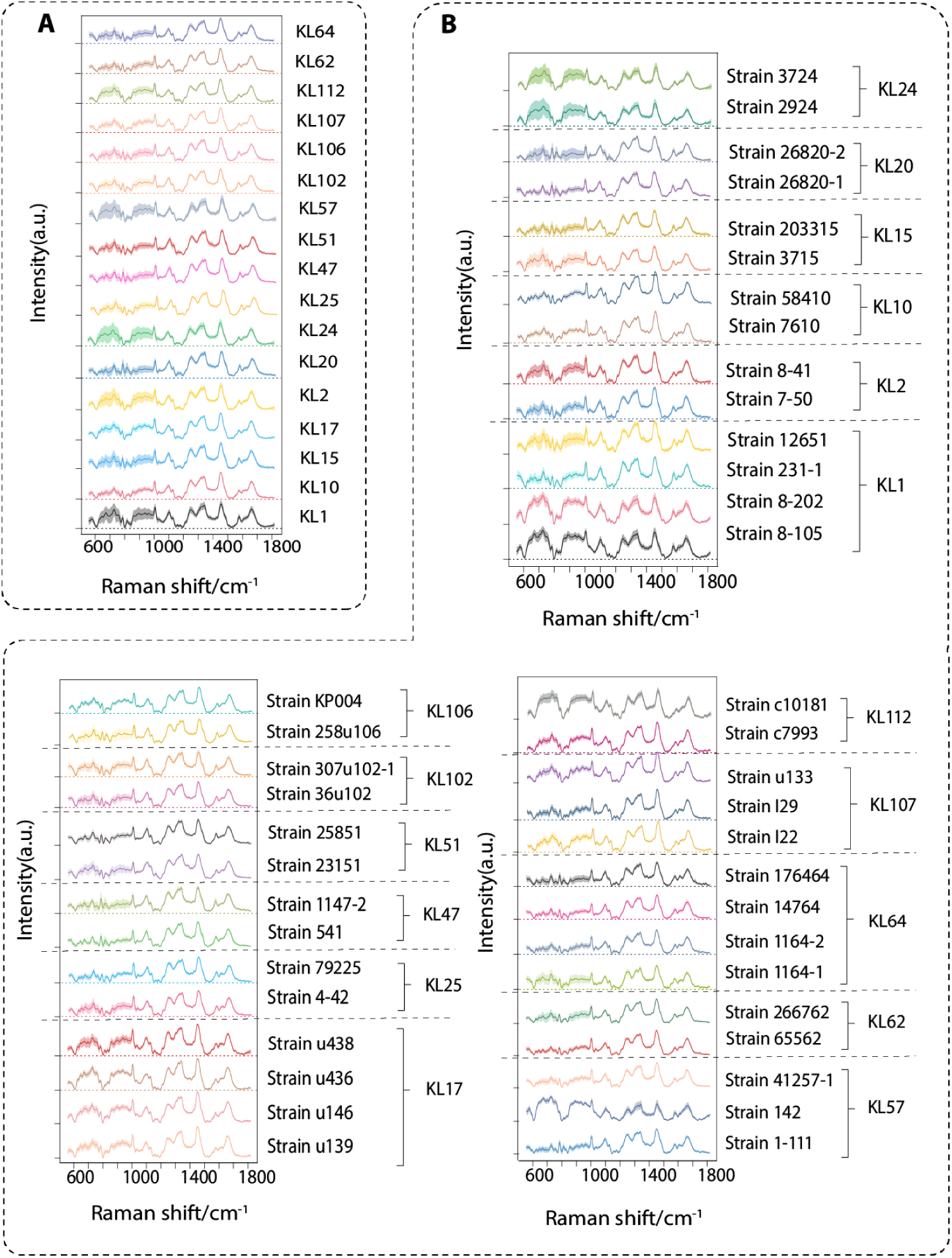
Average Raman spectra of 17 KL-types and corresponding 42 *K. pneumoniae* strains for AI model training. **(A)** Average Raman spectra of 17 KL-types: KL1, KL10, KL102, KL106, KL107, KL112, KL15, KL17, KL2, KL20, KL24, KL25, KL47, KL51, KL57, KL62, and KL64. **(B)** Average Raman spectra of 42 *K. pneumoniae* strains. The solid line represents the average Raman spectrum, and the shaded area represents the standard deviation.

### Development of Deep Learning Models for Phage Selection

Convolutional neural networks (CNN) have been widely applied to classify various microbial cells based on spectral data (*45-47*). However, we observed that traditional CNN performed poorly in distinguishing KL-types of MDR-hvKP strains. The extreme similarity of Raman spectra across strains likely limits the ability of standard CNNs to capture sufficient discriminatory features for accurate classification. To address this problem, we developed three enhanced CNN architectures: CNN_MLP, CNN_MLP-Attention, and CNN_MLP-Transformer. All these three models used a multilayer perceptron (MLP) to process the Raman spectral data and combined these with Raman image features. CNN_MLP-Attention and CNN_MLP-Transformer further incorporated attention mechanism and transformer encoders, respectively (Fig. 2).

**Fig. 2.**
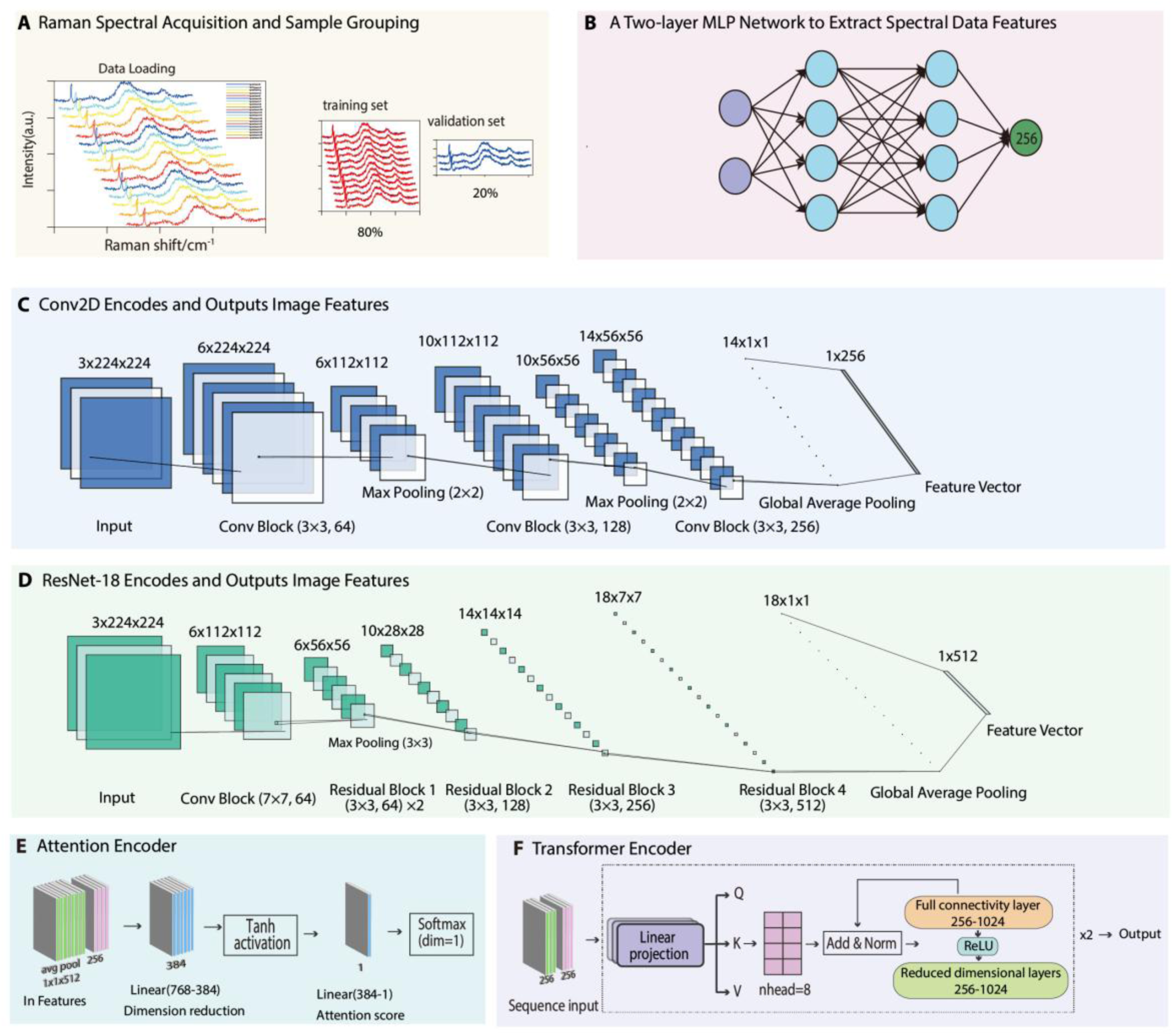
Structural frameworks of three deep leaning models. **(A)** Spectral data input and allocation into training and validation datasets. **(B)** Architecture of the two-layer multilayer perceptron (MLP) and extraction of spectral data features. **(C)** Conv2D-based encoding and extraction of spectral image features. **(D)** ResNet-18–based encoding and extraction of spectral image features. **(E)** Attention mechanism module of CNN_MLP-Attention model. **(F)** Transformer encoding module of CNN_MLP-Transformer model.

For training the CNN_MLP model, the Raman spectral data was randomly split into 80% as the training dataset and 20% as the validation dataset (Fig. 2A). In CNN_MLP, a two-layer MLP network was constructed to extract Raman spectral data features (Fig. 2B). In addition, we used a lightweight three-layer CNN to process the input Raman images and output 256-dimensional Raman image features (Fig. 2C). Finally, a fusion classifier connected the two branches through two linear layers to obtain the final classification outputs. As shown in Fig. 3A, CNN_MLP achieved an average KL-type prediction accuracy of 99.2%.

**Fig. 3.**
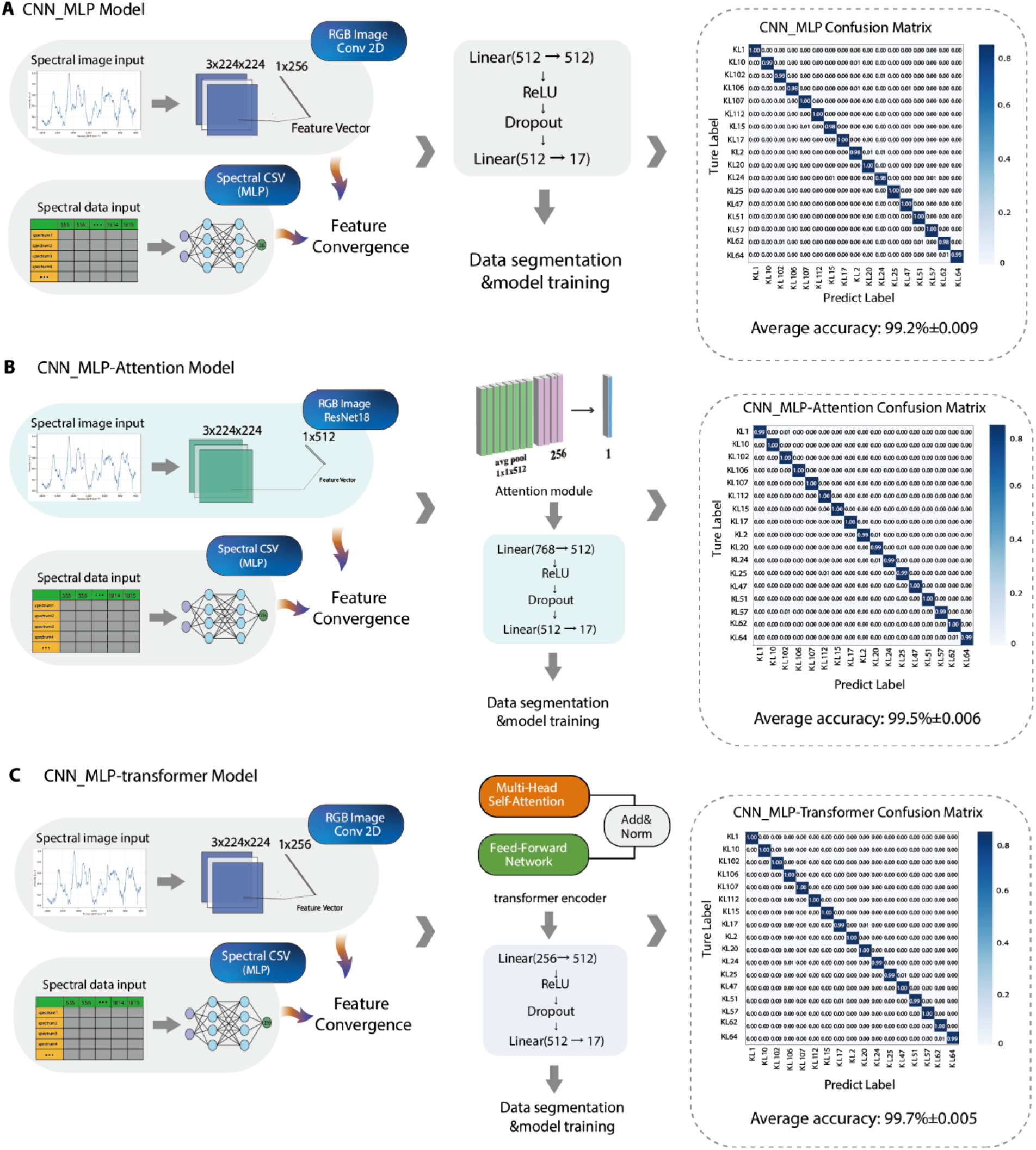
Architectures of three deep learning models and their corresponding confusion matrices for *K. pneumoniae* KL-type prediction. (A) CNN_MLP model. A three-layer lightweight CNN was used to extract 256-dimensional spectral image features, which were concatenated with 256-dimensional spectral features generated by an MLP. The fused feature vector was subsequently processed through two fully connected layers to produce the final classification output. The CNN_MLP model achieved an average classification accuracy of 99.2%. **(B)** CNN_MLP-Attention model. A ResNet-18 backbone was employed to extract 512-dimensional spectral image features, which were integrated with 256-dimensional MLP-derived spectral features. Following feature fusion via two fully connected layers, an attention module was applied to emphasize informative components within the fused representation, thereby refining the classification performance. This model attained an average accuracy of 99.5%. **(C)** CNN_MLP-Transformer model. A three-layer lightweight CNN and an MLP were used to extract 256-dimensional spectral image and spectral features respectively. After fusion through two fully connected layers, the resulting features were inputted into a two-layer transformer encoder for sequence-level feature modeling. The, transformer outputs were averaged along the sequence dimension to yield a unified 256-dimensional embedding, which was subsequently input into the classifier to generate the final prediction. This model achieved the highest average accuracy of 99.7%.

In CNN_MLP-Attention model, the lightweight three-layer CNN architecture was replaced by ResNet-18 to extract Raman image features (Fig. 2D). Unlike the CNN_MLP model, the CNN_MLP-Attention model outputted 512-dimensional Raman image features, while the extraction of Raman spectral data features employed the same MLP network as in the CNN_MLP model. The merging of these two modules produced 768-dimensional features. We applied an attention weighting module to learn assigning higher importance to key features in the fused vector (Fig. 2E). This attention-based fusion enabled the model to emphasize the interactions between the spectral data of interest and the image. With the same training-validation segmentation, the CNN_MLP-Attention model achieved average accuracy of 99.5% to distinguish 17 KL-types (Fig. 3B), demonstrating the benefits of attention-based weighting.

The transformer can effectively model long-range dependencies and global contextual relationships in Raman spectra, which traditional CNN alone may be hard to capture. Therefore, we incorporated a transformer encoder to facilitate deeper cross-modal interactions. A lightweight CNN was used as the image encoder to extract Raman image features, and an MLP network was used to extract Raman spectral data features (Fig. 2, B and C). The resulting image and spectral features were stacked along the modality dimension to form a feature sequence, which was then fed into a two-layer transformer encoder to model bidirectional interactions between the two modalities. Although the feature sequence length is limited, the transformer encoder provides a flexible attention-based mechanism to explicitly model interactions between spectral and image representations, enabling adaptive feature alignment and cross-modal fusion. After the transformer output, we averaged the sequence along the modality dimension to obtain the fused, unified feature representation (dimension 256), which was used for subsequent classification tasks. In this model construction, spectral data features were served as queries, while image features acted as keys and values in the transformer attention mechanism (Fig. 2F). This setup allowed the model to query the most relevant visual modalities in the image based on spectral cues, thereby facilitating more effective feature alignment and cross-modal fusion. The CNN_MLP-Transformer model achieved the highest accuracy of 99.7% (Fig. 3C). The confusion matrixes of three models are shown in Fig. 3, and the ROC curve evaluation results of the three models are presented in fig. S1.

### Model Validation using Independent Clinical Isolates

To independently evaluate model performance for selecting phages targeting an MDR-hvKP strain, we tested the three models on 10 uncharacterized clinical isolates which were identified as hvKP. For each isolate, Raman spectra were collected from approximately 100 individual cells per strain, and predictions for proper phages were generated using the three models.

All three models produced consistent phage type predictions for most isolates, with the exception of sample 6 (Fig. 4). The CNN_MLP model predicted multiple phage candidates for sample 6 with low confidence, that is 31% for KL15 phages, 22% for KL57 phages and lower probabilities for other KL-type phages (Fig. 4A). The CNN_MLP-Attention model also displayed uncertainty for sample 6, predicting a 45% probability for KL15 phages and 22% for KL57 phages (Fig. 4B). In contrast, the CNN_MLP-Transformer model predicted a 68% probability for KL15 and 8% for KL102 phages against sample 6, which showed a clearer dominance in candidate selection (Fig. 4C).

**Fig. 4.**
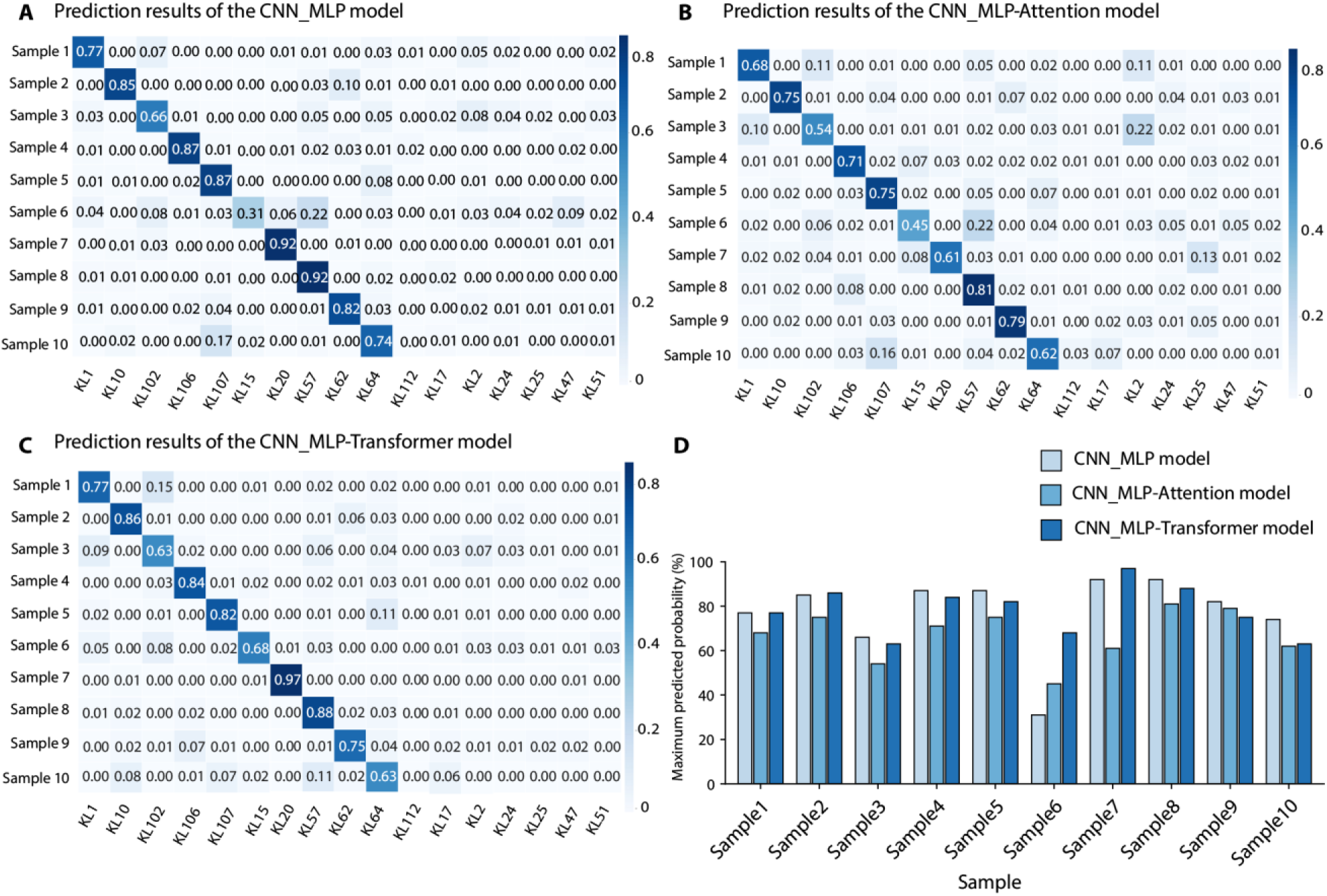
Confusion matrices for KL-type prediction of 10 uncharacterized hvKP clinical isolates using three deep learning models. **(A)** Predictions generated by the CNN_MLP model. **(B)** Predictions generated by the CNN_MLP-Attention model. **(C)** Predictions generated by the CNN_MLP-Transformer model. **(D)** Comparison of the maximum prediction outputs from the three models, showing substantial uncertainty in phage-type prediction for sample 6.

To verify these prediction results, we performed *in vitro* phage infection assays to determine whether the predicted phages were capable of lysing the corresponding clinical isolates. For each hvKP clinical isolate, phage lysis assays were conducted using the three highest-ranked KL-type-specific phages predicted by the CNN_MLP-Transformer model (Fig. 5A). The accession numbers of phages against a specific KL-types in our library are summarized in table S2. For eight clinical isolates (samples 1, 2, 3, 4, 5, 7, 8, and 10), the phage corresponding to the KL-type with the highest predicted probability successfully infected the bacteria and produced clear lysis plaques, while phages with lower predicted probabilities exhibited no detectable lytic activity (Fig. 5B and fig S2). For instance, sample 1 was predicted by the CNN_MLP-Transformer model to be of the KL1 type with a maximum probability of 77%. Consistent with this prediction, the KL1 phage RCIP0038 and RCIP0024 displayed pronounced lytic activity against sample 1, whereas KL102 (15% probability), KL57 (2% probability), and KL64 (2% probability) phages showed no infective activity (Fig. 5B). Interestingly, certain phages exhibit broad lytic activity. The KL10 phage RCIP0008 effectively lysed clinical isolates 2, 4, and 7 (Fig. 5B), indicating a broad host range spanning *K. pneumoniae* KL10, KL106, and KL20 types This broad host specificity warrants further investigation.

**Fig. 5.**
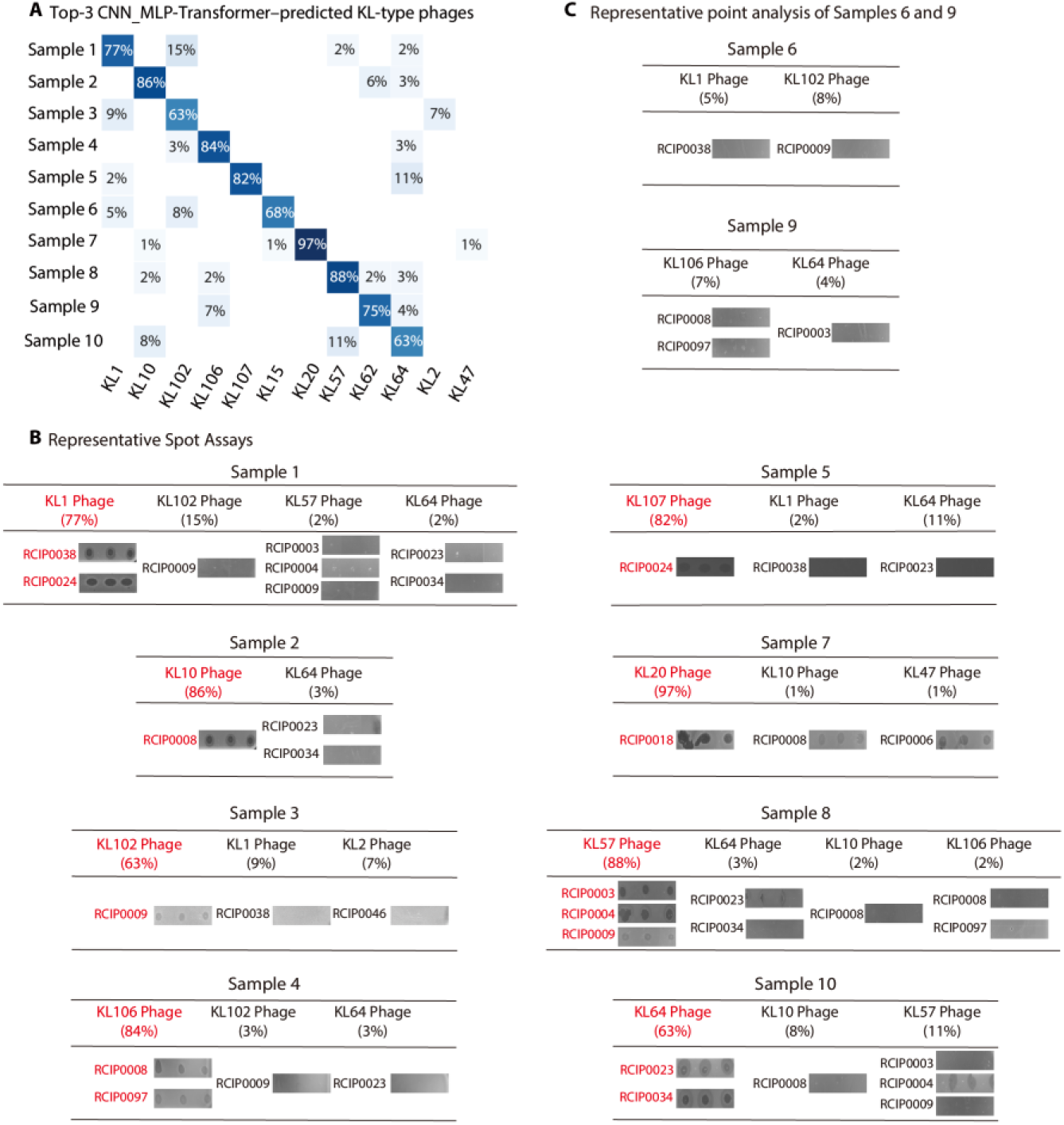
CNN_MLP-Transformer-guided phage infection detection. **(A)** The top three KL-types predicted by the CNN_MLP-Transformer model for each clinical isolate. **(B)** Representative results of phage lytic activity. Lysis was indicated by the formation of translucent plaques on bacterial culture medium. Plaque assays for each phage were performed in triplicate. Phages exhibiting effective lytic activity are highlighted in red, and the accession numbers of the phages used in the assays are indicated on the left. **(C)** Phage lysis assays for Samples 6 and 9. No phages corresponding to the highest-probability predicted KL-types of Samples 6 and 9 were available in our phage library. None of the lower-probability phages produced detectable translucent plaques.

The CNN_MLP-Transformer model predicted that samples 6 and 9 corresponded to KL15 (68% probability) and KL62 (75% probability), respectively. However, in our phage library, no phages capable of infecting KL15 or KL62 *K. pneumoniae* were available, precluding direct validation of these predictions. We therefore tested the phages with lower-ranked predictions, such as the KL102 phage (8% probability) for sample 6 and the KL106 phage (7% probability) for sample 9. However, neither exhibited detectable lytic activity (Fig. 5C).

Taken together, across the 10 clinical isolates, the CNN_MLP-Transformer model achieved an average phage selection accuracy of 78.3%. These findings demonstrate that the artificial intelligence-guided Raman spectroscopy enables rapid and relatively accurate phage selection at the single-cell level, thereby providing a foundation for personalized phage therapy against MDR-hvKP.

### Identification of key Raman Spectral features for accurate phage selection

Since Raman spectra reflect cellular biochemical composition, it is interest to investigate which spectral peaks are important for accurate phage selection. These peaks would facilitate us deeply understand the subtle biochemical differences among KL-types and biological basis for phage-bacterium interaction. To this end, an occlusion-based Raman spectra feature extraction method was employed to identify Raman peaks that are critical for accurate phage selection (*36*).

The feature extraction method identified several critical Raman peaks that are associated with specific biochemical components, providing insights into the molecular basis for phage selection. The identified peaks, such as the C-O-C glycosidic ring deformation (585 cm^-1^) and the O-P-O symmetric stretching of phosphates (971 cm^-1^), are highly indicative of the sugar-phosphate backbone of the capsular polymers. Furthermore, vibrations related to C-H/C-O-H bending (1375 cm^-1^) and ester groups (1745 cm^-1^) provide additional insights into the specific chemical modifications and potential lipid within the capsular structure (Table 1). These capsular-relevant spectral features are important for deciphering the biochemical basis of phage-bacterium interactions, as phages often recognize and bind to specific structures on the bacterial capsule.

**Table 1.**
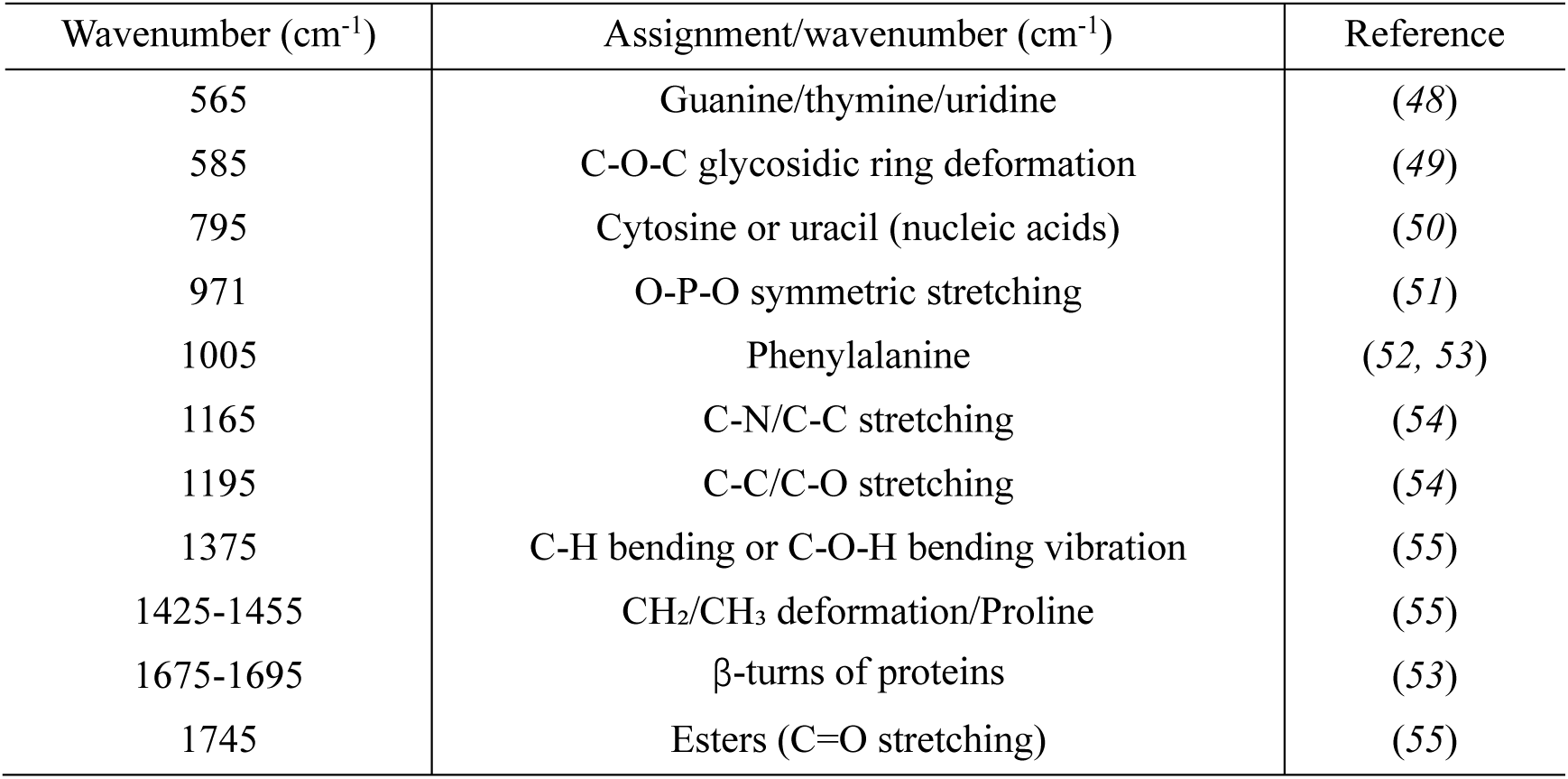
Characteristic Raman peak assignments of 17 KL-types of *K. pneumoniae*.

| Wavenumber ($\text{cm}^{-1}$) | Assignment/wavenumber ($\text{cm}^{-1}$) | Reference |
| --- | --- | --- |
| 565 | Guanine/thymine/uridine | (48) |
| 585 | C-O-C glycosidic ring deformation | (49) |
| 795 | Cytosine or uracil (nucleic acids) | (50) |
| 971 | O-P-O symmetric stretching | (51) |
| 1005 | Phenylalanine | (52, 53) |
| 1165 | C-N/C-C stretching | (54) |
| 1195 | C-C/C-O stretching | (54) |
| 1375 | C-H bending or C-O-H bending vibration | (55) |
| 1425-1455 | $\text{CH}_2/\text{CH}_3$ deformation/Proline | (55) |
| 1675-1695 | β-turns of proteins | (53) |
| 1745 | Esters (C=O stretching) | (55) |

By analyzing the spectral peak differences in these 17 KL-types *K. pneumoniae*, we observed that KL-types with similar biochemical profiles clustered together (Fig. 6). For instance, KL20 and KL102 were enriched for nucleic acid peaks (795 cm^-1^, 565 cm^-1^,). KL15 and KL51 showed prominent sugar related peaks (585 cm^-1^, 1375 cm^-1^). KL1, KL2, KL10, KL17, KL25, and KL57 collectively displayed strong protein-associated peaks (1005 cm^-1^, 1675-1695 cm^-1^, etc.). KL62, KL107, and KL112 were enriched for lipid peaks (1745 cm^-1^, 1195 cm^-1^, and 1165 cm^-1^), indicating that these three KL-types *K. pneumoniae* might have unique cell membrane or lipid metabolic features. These clustering patterns indicate possible spectral characteristics, metabolism and phenotype correlations that may warrant further exploration.

**Fig. 6.**
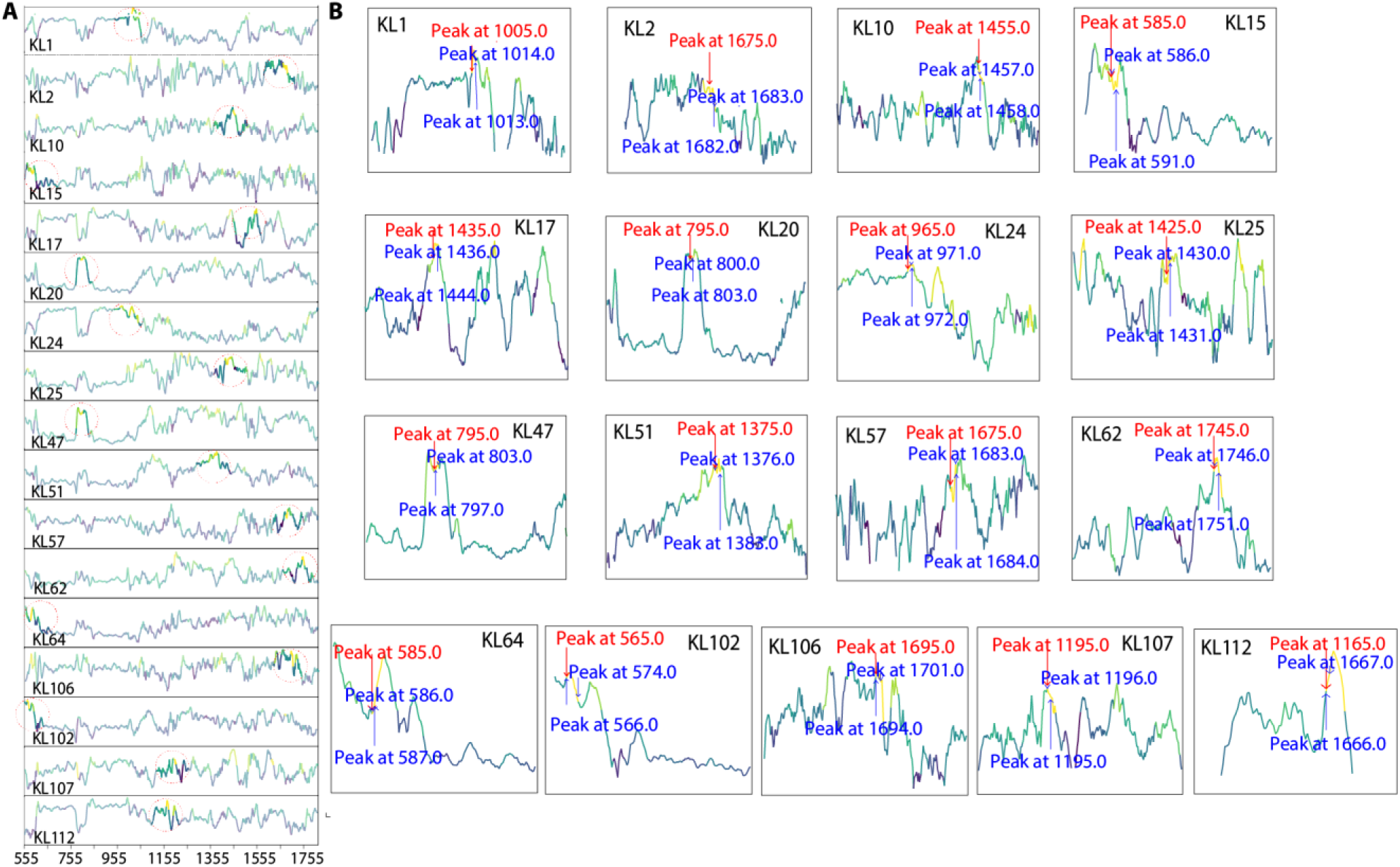
Characteristic Raman spectral peaks of 17 KL-types of *K. pneumoniae*. **(A)** Key peaks identified from the KL-type specific average Raman spectra through feature extraction, with the characteristic peak positions highlighted by red circle for each spectrum. **(B)** The top three peak positions derived from the average Raman spectral features of the 17 KL-types of *K. pneumoniae*.

## DISCUSSION

In this proof-of-concept study, we combined single-cell Raman spectroscopy with deep learning to enable rapid selection of lytic phages for multidrug-resistant hypervirulent *K. pneumoniae* (MDR-hvKP). By coupling single-cell Raman spectrum with deep learning, our workflow identified candidate therapeutic phages for clinical *K. pneumoniae* isolates with an average accuracy of 78.3%. This result highlights the practical potential of AI-guided Raman analysis to deliver time-efficient, strain-level phage matching—an essential requirement for precision phage therapy when conventional antibiotics fail.

Raman spectroscopy was adopted as the core analysis tool because its unique advantages that align with clinical needs. Raman spectroscopy is rapid, label-free, non-destructive, minimally affected by water interference, and adaptable to various pH, temperature, salty and oxygen content conditions. Meanwhile, when integration with laser tweezer, the laser tweezers Raman spectroscopy (LTRS) enables single cell analysis with minimal sample preparation, even in complex, polymicrobial specimens. This approach supports near-*in situ* analysis of individual bacterial cells in diverse clinical specimens such as blood, sputum, bronchoalveolar lavage fluid, urine, or cerebrospinal fluid, facilitating rapid, culture-free pathogen detection and phage selection. In addition, Raman spectroscopy provides rich biochemical information at the single-cell level, including detailed insights into biochemical components such as capsular polysaccharides and lipoproteins. These types of biochemical information are often critical determinants of phage-host specificity, making Raman spectra an informative basis for targeted phage matching.

Given the complexity and high dimensionality of single-cell Raman spectra, constructing a deep learning model capable of distinguishing subtle spectral differences among bacterial subtypes is essential. In this study, we designed and evaluated three deep learning architectures for lytic phage selection: conventional convolutional neural networks coupled to a multilayer perceptron (CNN_MLP), a CNN_MLP integrated with an attention module (CNN_MLP-Attention), and a CNN_MLP fused with a transformer encoder (CNN_MLP-Transformer). The CNN_MLP-Transformer architecture achieved the highest accuracy. It might leverage the transformer’s global attention mechanism, enhancing its ability to capture long-range feature dependencies in spectral data. While the CNN_MLP-Attention model improved local feature weighting relative to a conventional CNN, its capacity to model global spectral structures may be limited, resulting in slightly inferior performance in high-dimensional classification tasks.

To further interpret model decisions, we applied an occlusion-based feature analysis and identified Raman peaks that contributed most to accurate phage selection, facilitating partially decode the “black box” of artificial intelligence determination process. These subtle differences to influence phage binding among different bacterial subtypes in theory might be mapped to capsule polysaccharides, membrane proteins, and extracellular matrix.

The glycolic phosphate skeleton characteristics represented by the deformation of the C-O-C glycosidic ring and the symmetrical stretching of the O-P-O are specific signals of capsulated polysaccharides, while the stretching vibrations of C-N/C-C and C-C/C-O at 1165 cm^-1^ and 1195 cm^-1^ are commonly present in the extracellular matrix with complex components. These features manifest as subtle, distributed spectral shifts rather than isolated peak changes in the spectrum.

Such non-local spectral patterns are better captured by the transformer’s global attention, which likely aids discrimination of serotype-linked biochemical states relevant to phage receptor availability. By contrast, the attention module in CNN_MLP-Attention preferentially amplifies locally prominent peaks, making it less effective at integrating sparse, fingerprint-wide cues needed for robust subtype separation.

As a proof-of-concept, our result demonstrated the potential feasibility of Raman spectroscopy combined with deep learning for the selection of appropriate phages in clinical MDR-hvKP phage therapy. However, several critical steps are still required before this approach can be translated into clinical practice. First, a more comprehensive Raman spectral library of MDR-hvKP strains is necessitated. In this study, the spectral database included only 17 *K. pneumoniae* KL-types. In the future, we aim to expand this to cover all 158 KL-types. Greater breadth and depth will likely improve generalization and phage selection accuracy. Second, it is important to investigate the spectral differences between single bacterial cells derived from laboratory cultures and those directly obtained from patient specimens. To address this, a more robust deep learning model must be developed to handle potential subtle variations among different domains. Such advancements will facilitate the extension of laboratory-based spectral libraries to real-world clinical applications. Third, the establishment of a more complete therapeutic phage library will further support clinical implementation. This requires extensive efforts in the isolation and characterization of novel phage candidates. In parallel, comprehensive multi-omics analyses, including genomic, transcriptomic, proteomic, and Raman spectral profiling at the single-cell level, will deepen our understanding of the complex interactions between bacteria and phages. These insights will ultimately enhance the precision and efficacy of future phage therapy strategies.

## Materials and Methods

### Strains and Cultivation

Forty-two multidrug-resistant hypervirulent *K. pneumoniae* strains representing 17 KL-types which account for the majority of MDR-hvKP infections globally were selected(*28, 44*). The complete list of these strains is provided in table S1. All strains were cultured in Luria Bertani (LB) medium (10 gL^-1^ tryptone, 5 gL^-1^ yeast extract, and 10 gL^-1^ NaCl) and incubated at 37 °C. In addition, 10 clinical isolates were taken from the clinical laboratory and cultured under the same conditions before Raman measurement. Bacteriophage particles were precipitated with 10% (w/v) polyethylene glycol 8000 and 0.5 M NaCl at 4°C for 24 hours to obtain the purified phage solution(*56*).

### Single-cell Raman Spectroscopy and Data Processing

Single-cell Raman spectra were acquired using a Laser Tweezers Raman Spectroscopy (LTRS) system as previously described(*36, 41*). Briefly, 1 mL of bacterial culture in the logarithmic growth phase was centrifuged and washed three times with 0.9% sodium chloride (NaCl) solution. The resulting cell pellet was resuspended in 1 mL of 0.9% NaCl. A 100 μL aliquot of bacterial suspension was then transferred onto a quartz chip within a sealed biosafety-grade miniature chamber. Prior to acquiring the bacterial Raman spectra, the LTRS system was calibrated using the characteristic Raman peaks of polystyrene microspheres at 620.9 cm^-1^, 1001.4 cm^-1^, and 1602.3 cm^-1^. The integration time for each single-cell measurement was set to 25 seconds.

Raman spectral data was first converted to ASCII files using the Winspecs software and subsequently processed with the "Ramanpro 0.4.2" software package that was developed in the R language. The spectra then underwent cosmic ray removal and background subtraction. Smoothing was performed using a Savitzky-Golay filter, and fluorescence background was eliminated via polynomial baseline fitting. Finally, the spectral data were normalized using vector normalization.

### Deep Learning Model Training and Evaluation

Three multimodal deep learning architectures were developed to explore the fusion of image and spectral data: CNN_MLP, CNN_MLP-Attention, and CNN_MLP-Transformer. The CNN_MLP serves as a baseline model, integrating convolutional and multilayer perceptron networks. In all architectures, image features extracted by CNNs (specifically, ResNet-18 was used in CNN_MLP-Attention) were fused with spectral features processed by an MLP. The CNN_MLP-Attention model further incorporates attention mechanisms, while the CNN_MLP-Transformer employs a transformer encoder to enhance cross-modal representation learning.

In the model training, the spectral data was split 80%/20% into training and validation dataset. The Multimodal Model, which combined image and spectral features, was initialized with a preset training round of 100 times. The model was trained iteratively within a specified epoch: the image, spectral data, and labels were moved to the GPU in each batch, and forward propagation was performed to calculate the loss, while backward propagation was used to update the weights. After each epoch, the accuracy of the training set was synchronously calculated, and gradient free evaluation was performed on the validation dataset to calculate the same metrics.

We applied the same preprocessing process as the training data to 10 sets of validation to test the three constructed classification models. The prediction results showed the prediction labels and corresponding KL-types of each set of data. The entire process strictly maintains consistency with data processing during the training stage to ensure the reliability of the assessment.

### Bacteriophage selection and Plaque Assays

To validate the model’s performance in phage selection against clinical strains, ten hvKP clinical isolates were cultured. Following culture, approximately 100 single cells were randomly selected from each isolate, and Raman spectra were obtained. According to the predictions of KL-types by our deep learning models, the top three KL-type phages were selected from our phage library for infection assays against the clinical isolates.

The lytic activity of the selected phages was assessed using the double-layer agar plate method as previously described. Briefly, exponentially growing cultures of the clinical isolates were mixed with soft agar and overlaid onto a base plate to form a lawn. Following overlay preparation, each phage lysate (1×10^9^ PFU/mL, 3 µL) was spotted onto the corresponding bacterial lawn. After the overlay solidified, the formation of clear lysis plaques at the spotting sites, indicated successful phage-mediated lysis, or non-lytic in the absence of visible plaques. Each phage-isolate combination was tested in triplicate to ensure reproducibility(*57*).

## Supplementary Materials

**This PDF file includes:**

Figs. S1 to S2

Tables S1 to S2

## Acknowledgments

We thank Dr Hui Wang, Department of Clinical Laboratory, Peking University People’s Hospital, Beijing, China, and Dr Qiwen Yang, Peking Union Medical College Hospital, Beijing, China, for their provision of the microbial resources.

## Author contributions

Conceptualization: J.F., C.W., J.W., Y.V.F. Methodology: B.L., C.W., W.L., J.W., X.G. Investigation: B.L., C.W., Q.Z., W.L. Visualization: B.L., C.W., Q.Z., J.W. Supervision: J.W., J.F., Y.V.F. Writing—original draft: B.L., Q.Z., Y.V.F. Writing—review & editing: B.L., C.W., Q.Z., J.W., W.L., J.F., Y.V.F., X.G.

## Data and materials availability

All data are available in the main text or the supplementary materials.

