## Supplementary for "Single-Cell Raman Profiling Enables Rapid Precision Phage Therapy Against Multidrug-Resistant Hypervirulent *Klebsiella pneumoniae*"

Science Advances
AAAS

### Supplementary Materials for

#### Single-Cell Raman Profiling Enables Rapid Precision Phage Therapy Against Multidrug-Resistant Hypervirulent *Klebsiella pneumoniae*

Beimin Liu *et al.*

**This PDF file includes:**

Figs. S1 to S2
Tables S1 to S2

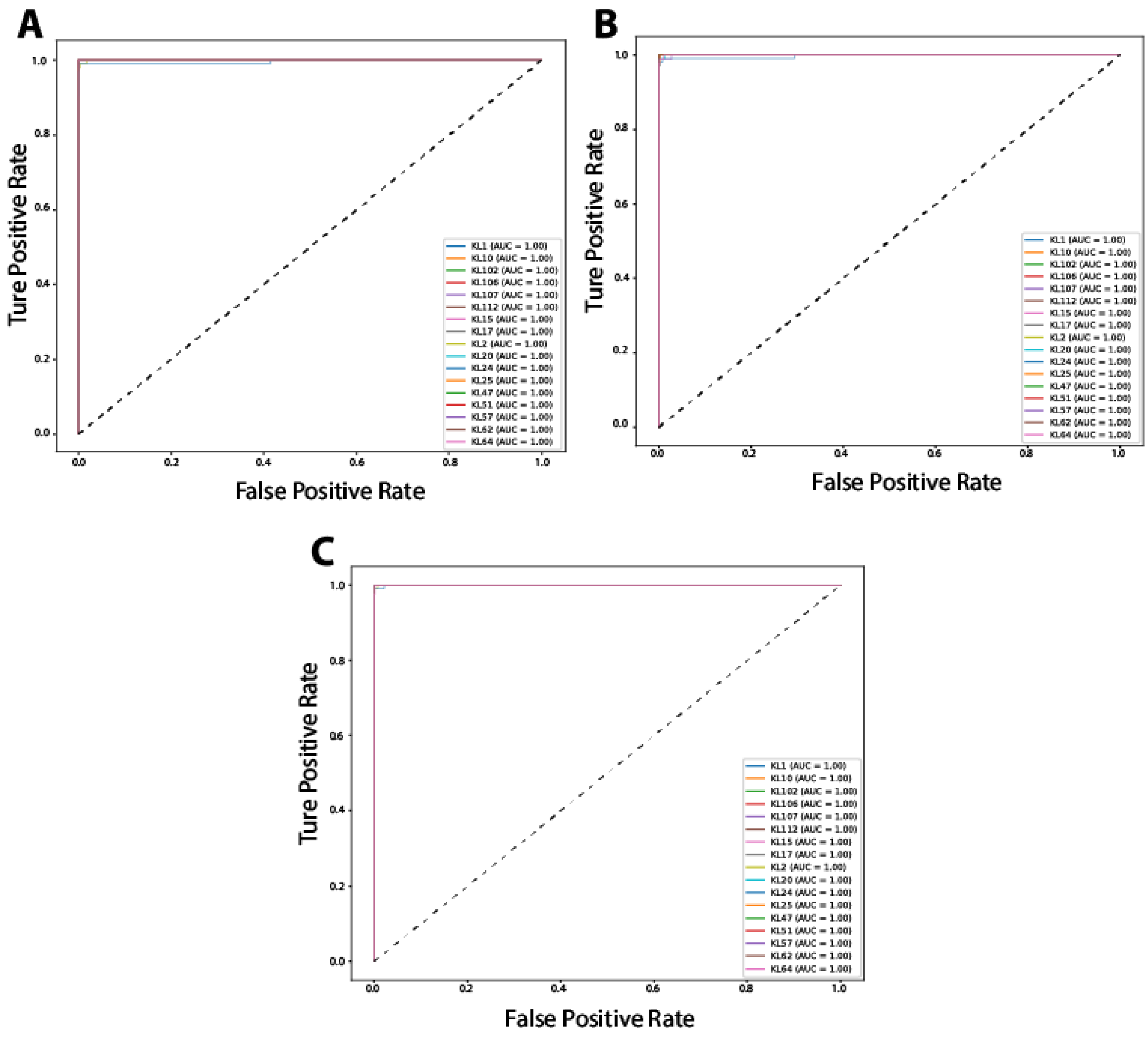


**Figure S1. Multi-class receiver operating characteristic (ROC) curves for the classification of KL-types using three deep learning models.** ROC curves for each KL-type were generated using **(A)** CNN_MLP, **(B)** CNN_MLP-Attention, and **(C)** CNN_MLP-Transformer. All three models demonstrated near-perfect classification performance across KL-types, with area under the curve (AUC) values approaching 1.00, indicating strong discriminative capability and robustness of the proposed architectures.

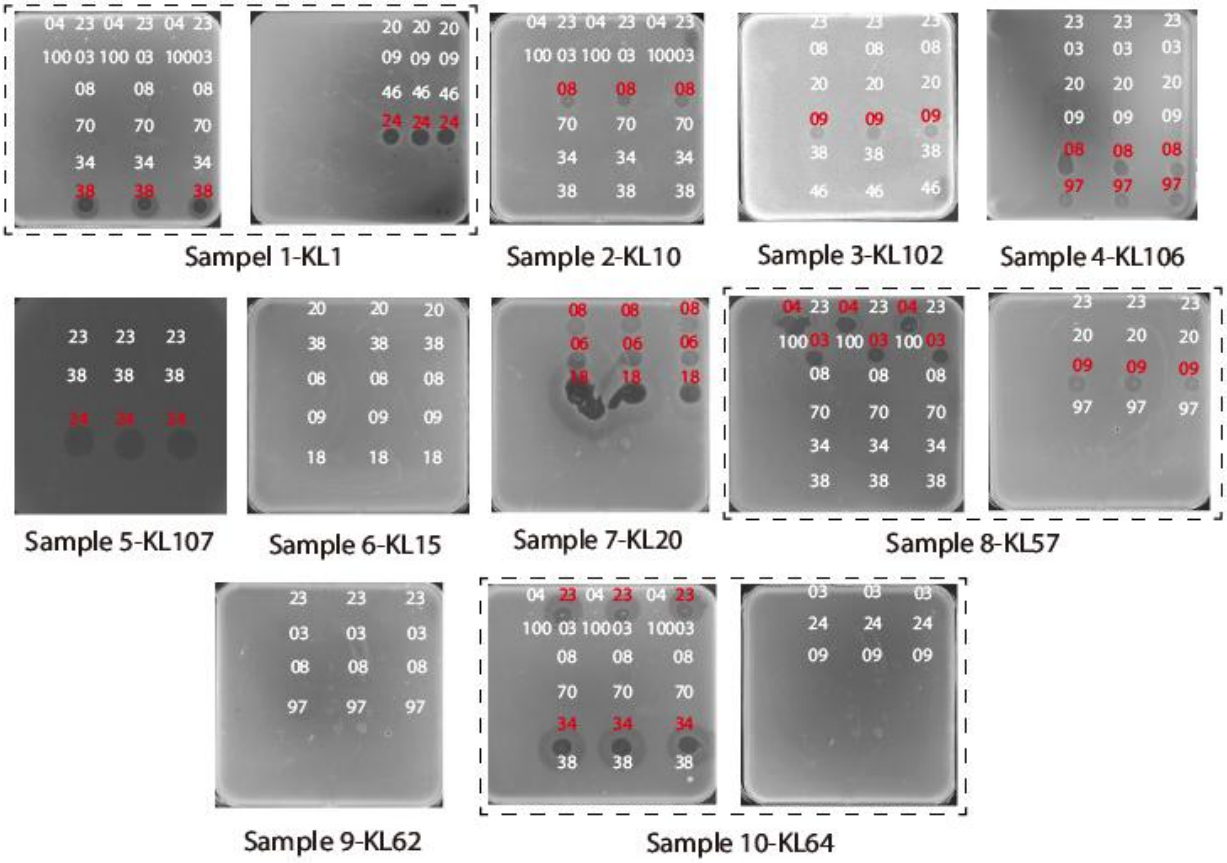


**Figure S2. Experimental validation of phage lytic activity against 10 uncharacterized hypervirulent *K. pneumoniae* clinical isolates.** Representative agar plate images show the lysis outcomes of phages tested against 10 uncharacterized hypervirulent *K. pneumoniae* clinical isolates. For each isolate, phage lysis assays were performed using the phage corresponding to the KL-type with the highest predicted probability generated by the CNN_MLP-Transformer model. Each phage-isolate combination was tested in triplicate, and phage accession numbers were indicated on the agar surface. Clear plaque formation indicates effective bacterial lysis, and phages exhibiting positive lytic activity are highlighted in red. For example, Sample 1 was predicted to be of the KL1 type. Consistent with this prediction, the KL1 phage RCIP0038 (abbreviated as 38) and RCIP0024 (abbreviated as 24) produced clear lysis plaques on Sample 1, confirming their lytic activity and were therefore marked in red. Notably, no phages capable of infecting KL15 or KL62 *K. pneumoniae* strains were available in our phage library, precluding direct experimental validation of these predictions.

**Table S1. Fifty multidrug-resistant hypervirulent *K. pneumoniae* strains representing 17 KL-types**

| KL-types and strains | | Number of spectra |
|---|---|---|
| KL1 | Strain 12651<br>Strain 231-1<br>Strain 8-202<br>Strain 8-105 | 404 |
| KL10 | Strain 58410<br>Strain 7610 | 353 |
| KL102 | Strain 307u102-1<br>Strain 36u102 | 354 |
| KL106 | Strain KP004<br>Strain 258u106 | 358 |
| KL107 | Strain u133<br>Strain I29<br>Strain I22 | 400 |
| KL112 | Strain c10181<br>Strain c7993 | 341 |
| KL15 | Strain 203315<br>Strain 3715 | 346 |
| KL17 | Strain u438<br>Strain u436<br>Strain u146<br>Strain u139 | 406 |
| KL2 | Strain 8-41<br>Strain 7-50 | 346 |
| KL20 | Strain 26820-2<br>Strain 26820-1 | 356 |
| KL24 | Strain 3724<br>Strain 2924 | 358 |
| KL25 | Strain 79225<br>Strain 4-42 | 344 |

| | | |
|---|---|---|
| KL47 | Strain 1147-2<br>Strain 541 | 344 |
| KL51 | Strain 25851<br>Strain 23151 | 350 |
| KL57 | Strain 41257-1<br>Strain 142<br>Strain 1-111 | 418 |
| KL62 | Strain 266762<br>Strain 65562 | 358 |
| KL64 | Strain 176464<br>Strain 14764<br>Strain 1164-2<br>Strain 1164-1 | 418 |

**Table S2. Phages with accession numbers confirmed to exhibit lytic activity against *K. pneumoniae* KL-types in our phage library**

| KL-type | Phage | Accession No. |
|---|---|---|
| KL1 | RCIP0038 | NMDC60139417 |
| | RCIP0024 | NMDC60139403 |
| KL10 | RCIP0008 | NMDC60139387 |
| KL102 | RCIP0009 | NMDC60139388 |
| KL106 | RCIP0008 | NMDC60139387 |
| | RCIP0097 | NMDC60139476 |
| KL107 | RCIP0024 | NMDC60139403 |
| KL2 | RCIP0046 | NMDC60139425 |
| KL20 | RCIP0008 | NMDC60139387 |
| | RCIP0006 | NMDC60139385 |
| | RCIP0018 | NMDC60139397 |
| KL47 | RCIP0006 | NMDC60139385 |
| KL57 | RCIP0003 | NMDC60139382 |
| | RCIP0004 | NMDC60139383 |
| | RCIP0009 | NMDC60139388 |
| KL64 | RCIP0023 | NMDC60139402 |
| | RCIP0034 | NMDC60139413 |
